## Supplementary file 1 for "3D directional tuning in the orofacial sensorimotor cortex during natural feeding and drinking"

### Supplementary figures and tables

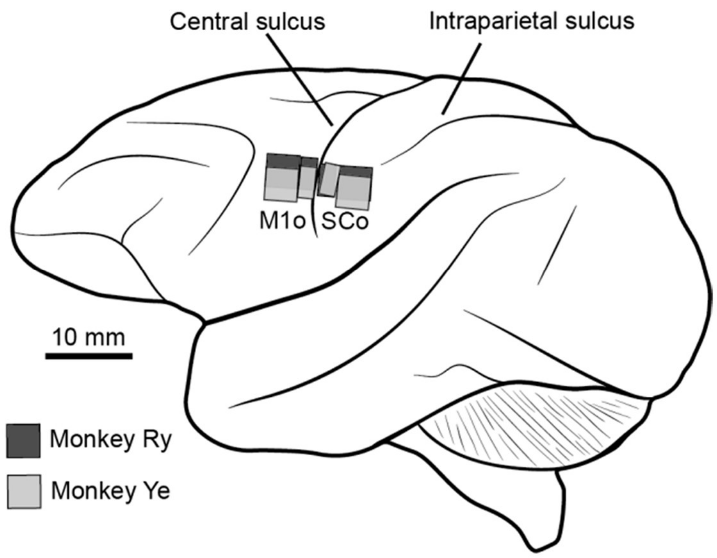


***Supplementary file 1 - Figure 1.*** *Microelectrode array locations. Squares represent Utah arrays and rectangles represent floating microelectrode arrays. Drawing is to scale and array locations were taken from surgical photographs. Adapted from Supplementary Figure 8 of "Robust cortical encoding of 3D tongue shape during feeding in macaques" (Laurence-Chasen et al., 2023).*

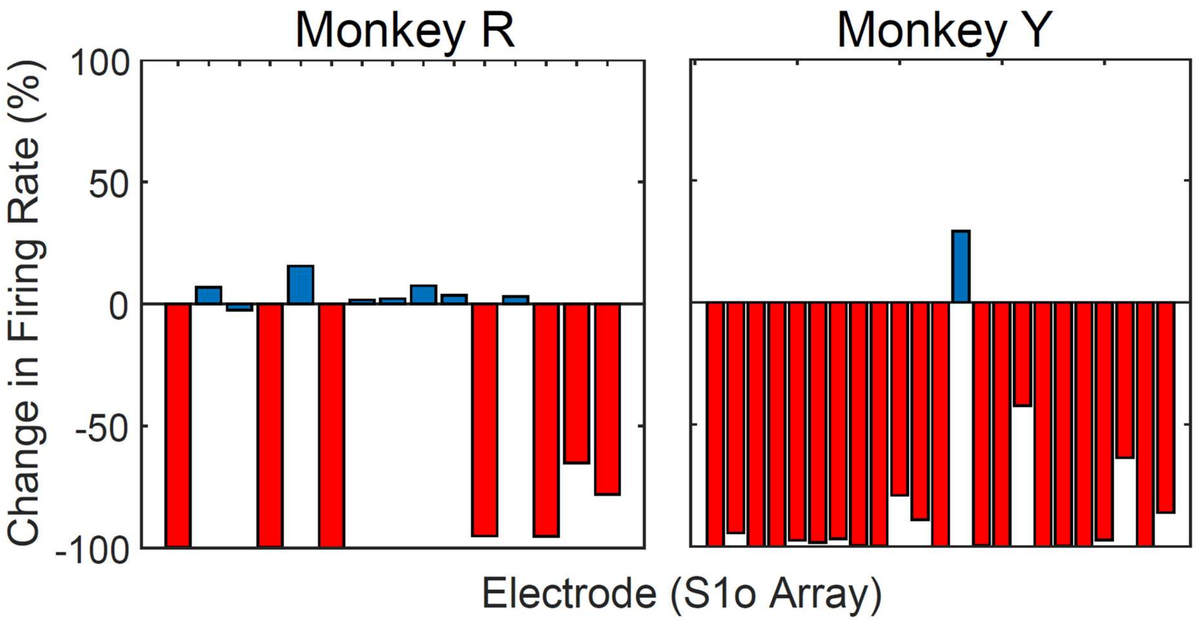


*__Supplementary file 1 – Figure 2.__ Change in firing rates of cortical somatosensory neurons. A decrease in firing rate in over 50% of SIo neurons was the criterion for a successful nerve block. Red bars indicate a relative decrease in firing rate for a given electrode, while blue bars indicate an increase. Only electrodes for which the control baseline firing rate was greater than 3 spikes/s are shown. Adapted from Supplementary Figure 6 of "Loss of oral sensation impairs feeding performance and consistency of tongue–jaw coordination" (Laurence-Chasen et al., 2022).*

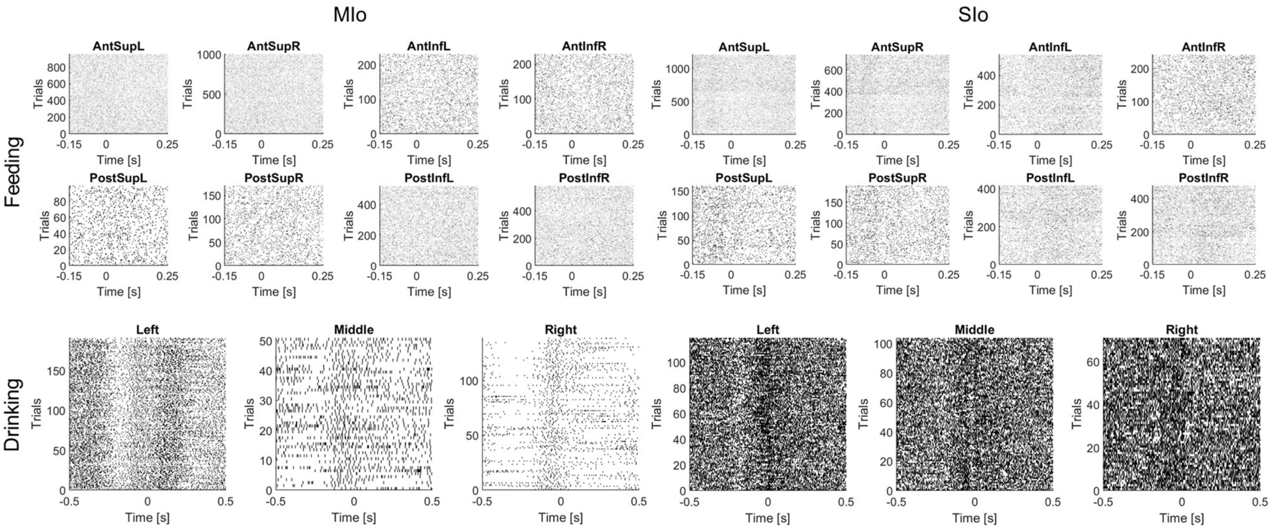


*__Figure 2 – figure supplement 1.__ Directional neural responses across trials. Raster plots corresponding to the same neurons represented in Figure 2. All trials in the respective dataset are included, grouped by direction.*

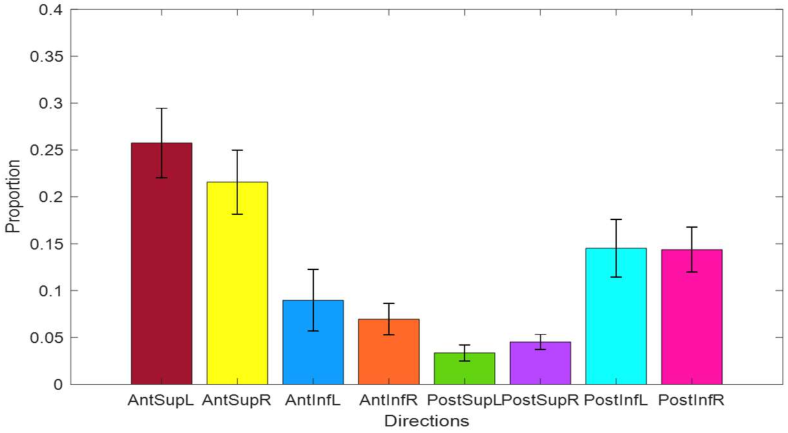


*Figure 3 – figure supplement 1*. Proportion of feeding trials in each group of directions. Error bars represent ±1 standard deviation across datasets (n = 4).

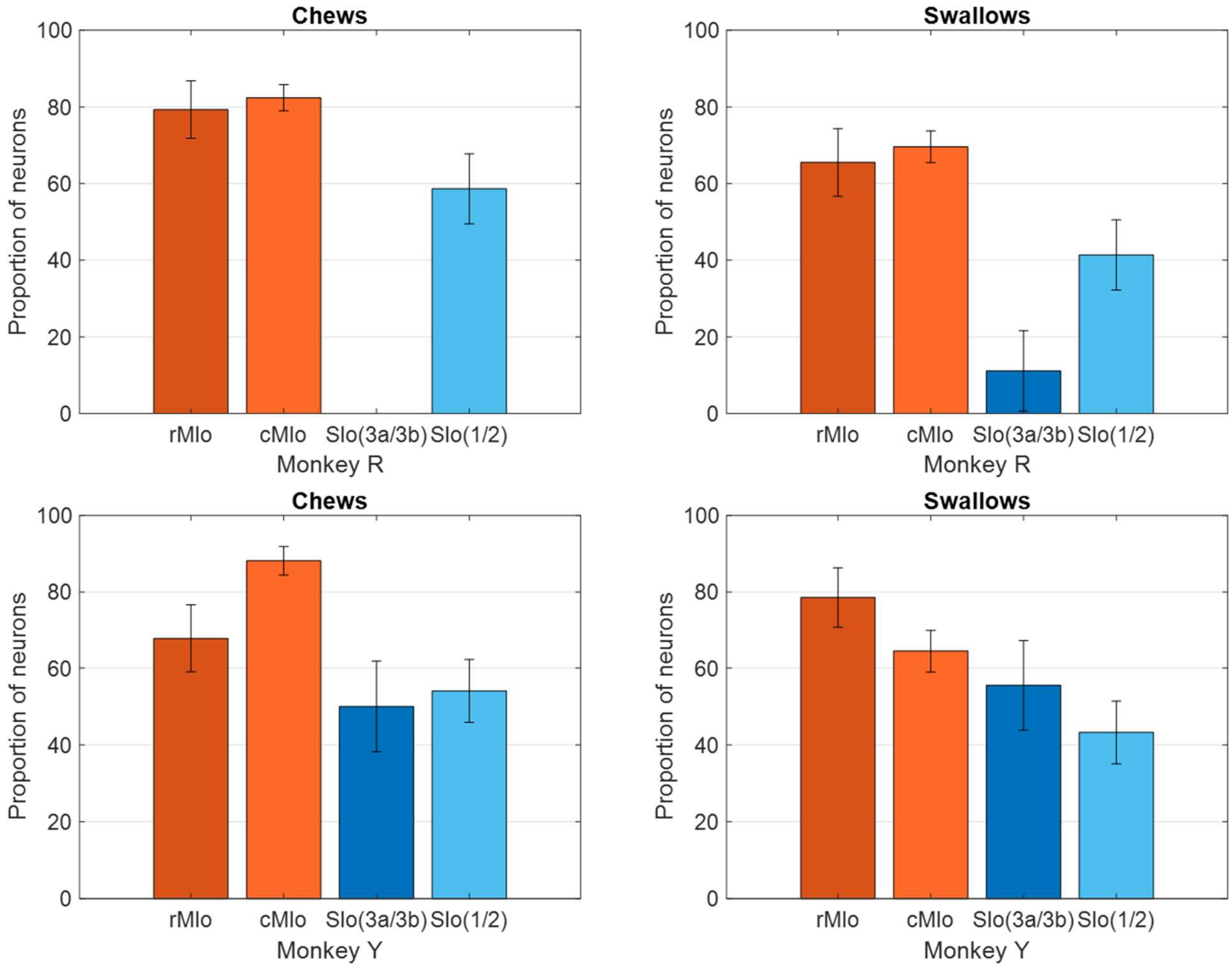


*Figure 3 – figure supplement 2.* Proportion of neurons directionally modulated during chews versus swallows in both monkeys. Recordings were taken from four areas of the OSMCx: rMIo - rostral M1, cMIo - caudal M1, SIo(3a/3b) - area 3a/3b, and SIo(1/2) - area 1/2. Error bars represent ±1 SE.

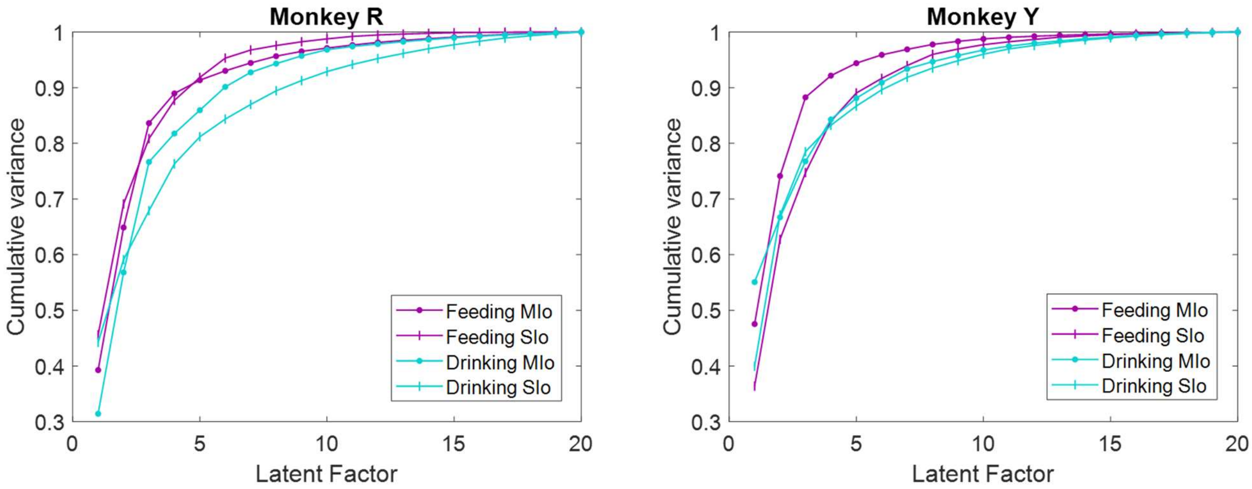


***Figure 7 – figure supplement 1***. *Comparison of cumulative explained variance between feeding and drinking behaviors with an equal number of neurons (N = 24), for both subjects.*

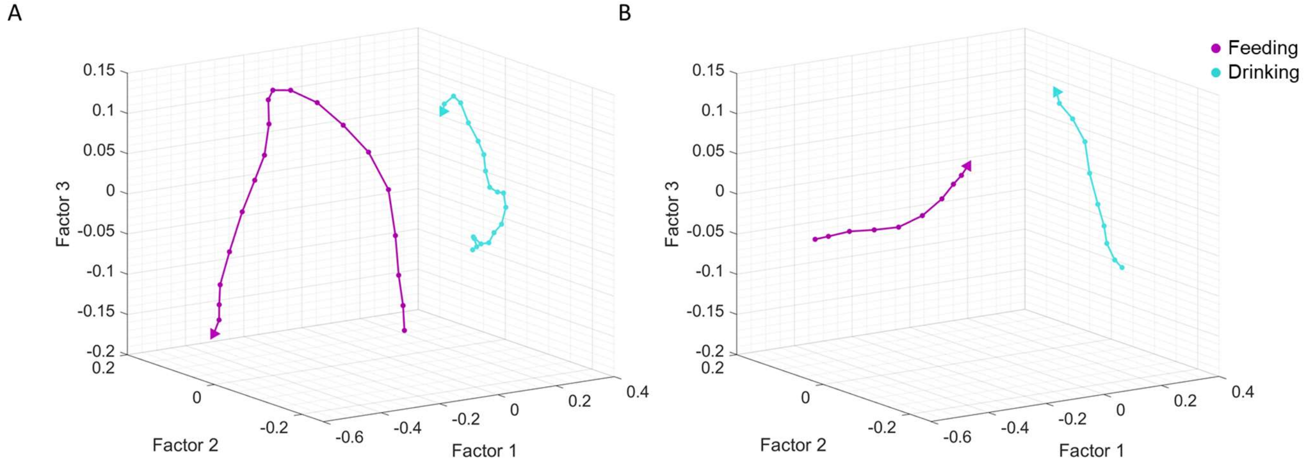


***Figure 7 – figure supplement 2***. *Comparison between stable MIo (N = 20) neural population trajectories in feeding and drinking behaviors for* ***(A)*** *all trials 200 ms around minimum gape and* ***(B)*** *a subset of trials (feeding: N = 40, drinking: N = 175) with similar kinematics 100 ms after minimum tongue protrusion. Dots represent each 10 ms time bin and arrow represents the end of the trajectory. Data from monkey Ry.*

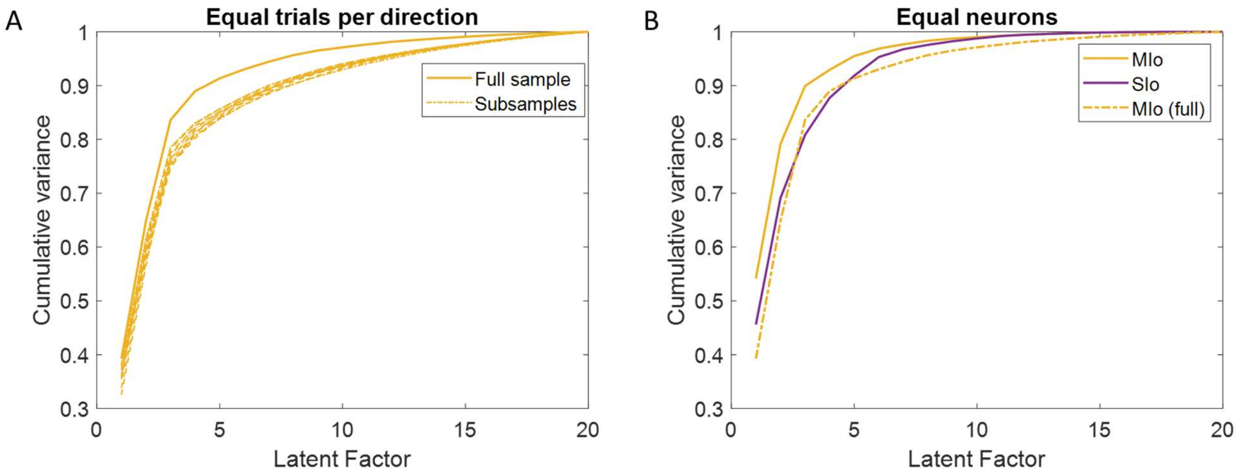


***Figure 7 – figure supplement 3****. Effect of subsampling* ***(A)*** *equal number of trials per direction (N = 80) from MIo and* ***(B)*** *equivalent neuron counts from MIo and SIo populations (N = 24) on the cumulative variance explained by latent factors.*

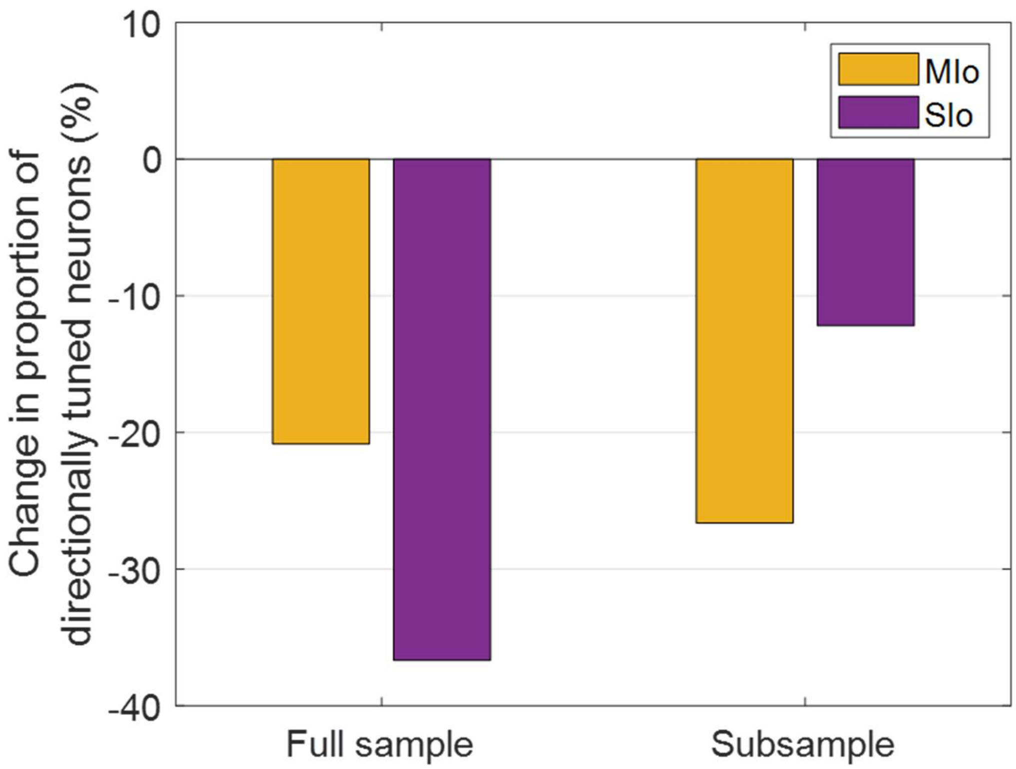


***Figure 10 – figure supplement 1.*** *Effect of subsampling drinking trials with similar kinematic profiles on the change in proportion of directionally tuned neurons in control vs. nerve block conditions for both subjects.*

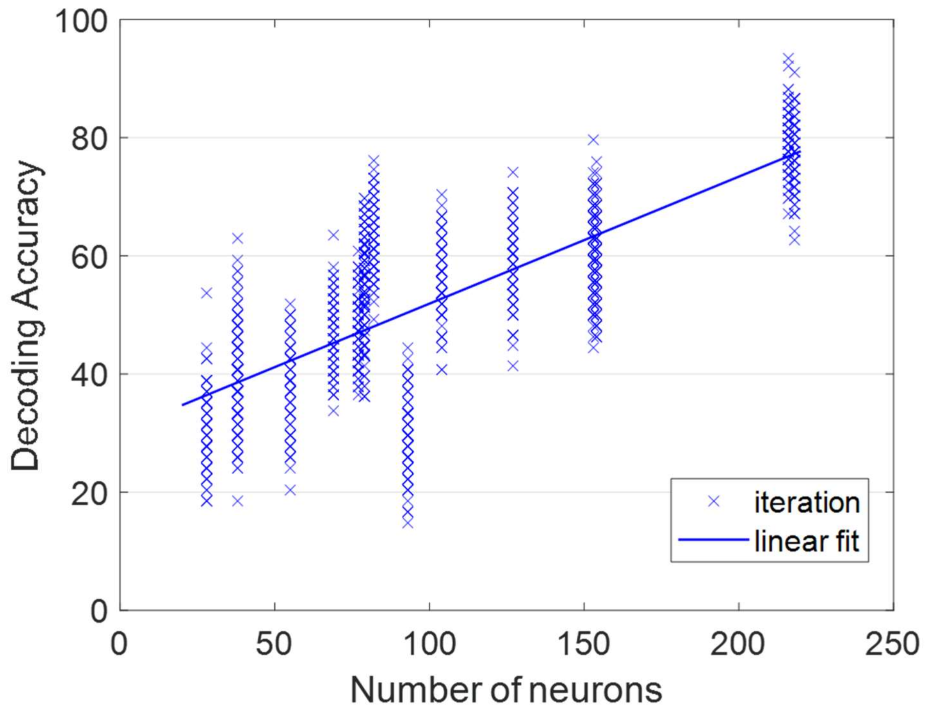


*Figure 13 – figure supplement 1.* Correlation between population size and decoding accuracy. Each cross represents one iteration of KNN classification, and the trendline is the linear fit: $R^2 = 0.6$.

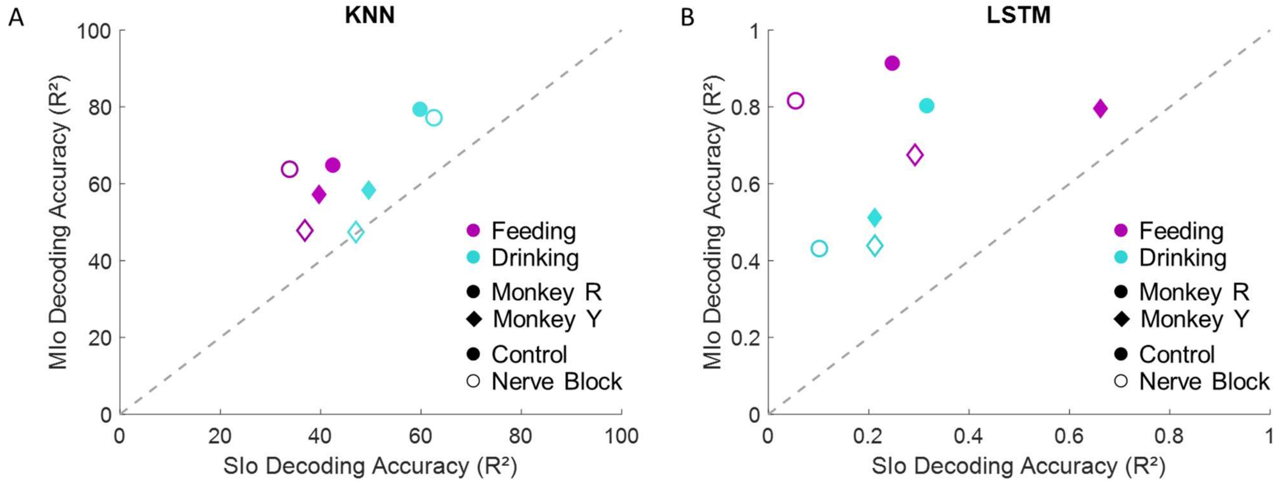


*Figure 13 – figure supplement 2.* Decoding accuracies from neuronal populations of various sizes. ***(A)*** Comparison between average decoding accuracy of KNN classifier. Chance level is 33.33%. ***(B)*** Comparison between average decoding accuracy by LSTM network. Data shown separately for each subject, behavioral task, and condition. The dashed line signifies equal decoding performance for MIo and SIo.

| Control | rMIo | cMIo | SIo(3a/3b) | SIo(1/2) |
|---|---|---|---|---|
| Ry Feeding | 29 | 125 | 9 | 29 |
| Ye Feeding | 28 | 76 | 18 | 37 |
| Ry Drinking | 31 | 185 | 23 | 54 |
| Ye Drinking | 23 | 104 | 16 | 63 |
| **Nerve Block** | | | | |
| Ry Feeding | 27 | 126 | 1 | 27 |
| Ye Feeding | 29 | 64 | 17 | 21 |
| Ry Drinking | 36 | 182 | 26 | 56 |
| Ye Drinking | 22 | 55 | 14 | 55 |

***Supplementary file 1 - Table 1.*** *Numbers of individual neurons recorded from each array location during each data collection session.*
