## Supplementary file 2 for "3D directional tuning in the orofacial sensorimotor cortex during natural feeding and drinking"

### Analysis of lingual yaw and pitch during feeding

To get a better understanding of directional tuning in three dimensions, we analyzed the lateral and vertical components of 3D tongue direction independently. The laterality of movements (yaw) was determined as the angle of rotation about the vertical axis (α) and the vertical component (pitch) was the angle of rotation about the horizontal plane (ϕ):

$$\alpha = \tan^{-1}(\Delta z/\Delta x) \quad (1)$$

$$\varphi = \tan^{-1}\Delta y/\sqrt{\Delta x^2 + \Delta z^2} \quad (2)$$

Where $x, y, z$ represent the position along the Posterior-Anterior, Inferior-Superior, and Left-Right axes respectively and $\Delta x, \Delta y, \Delta z$ represent the directional change in the position of the tongue tip over each 100 ms interval.

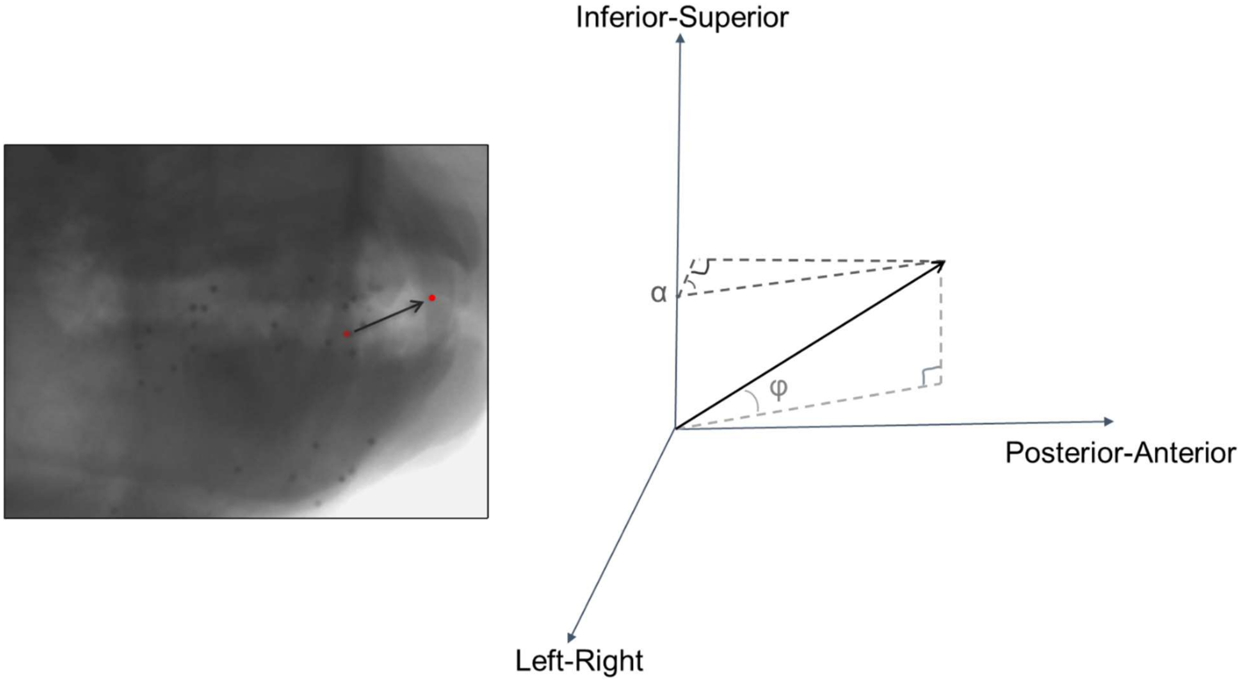


***Supplementary file 2 - Figure 1.*** *Calculation of directional angles. The vector represents the direction of movement over a 100 ms interval, where α describes the angle of lateral rotation (yaw) and φ describes the angle of the change in elevation (pitch) of the tongue tip.*

These results can be interpreted as the 3D direction of tongue movement broken down into its components. Generally, we made similar observations across the 3D angle, yaw, and pitch. This serves to reinforce our main results about the modulation of the OSMCx to tongue direction during feeding and how it is affected by the loss of oral tactile sensation.

**Distribution of yaw and pitch during feeding**

There were similar distributions of yaw and pitch observed in both monkeys. For yaw, there was a multimodal distribution, with three peaks in the far right (positive), center, and far left. Conversely, there was a somewhat uniform distribution for pitch in the control condition, with the center near 0°, which became skewed toward the upward (positive) directions under the nerve block condition.

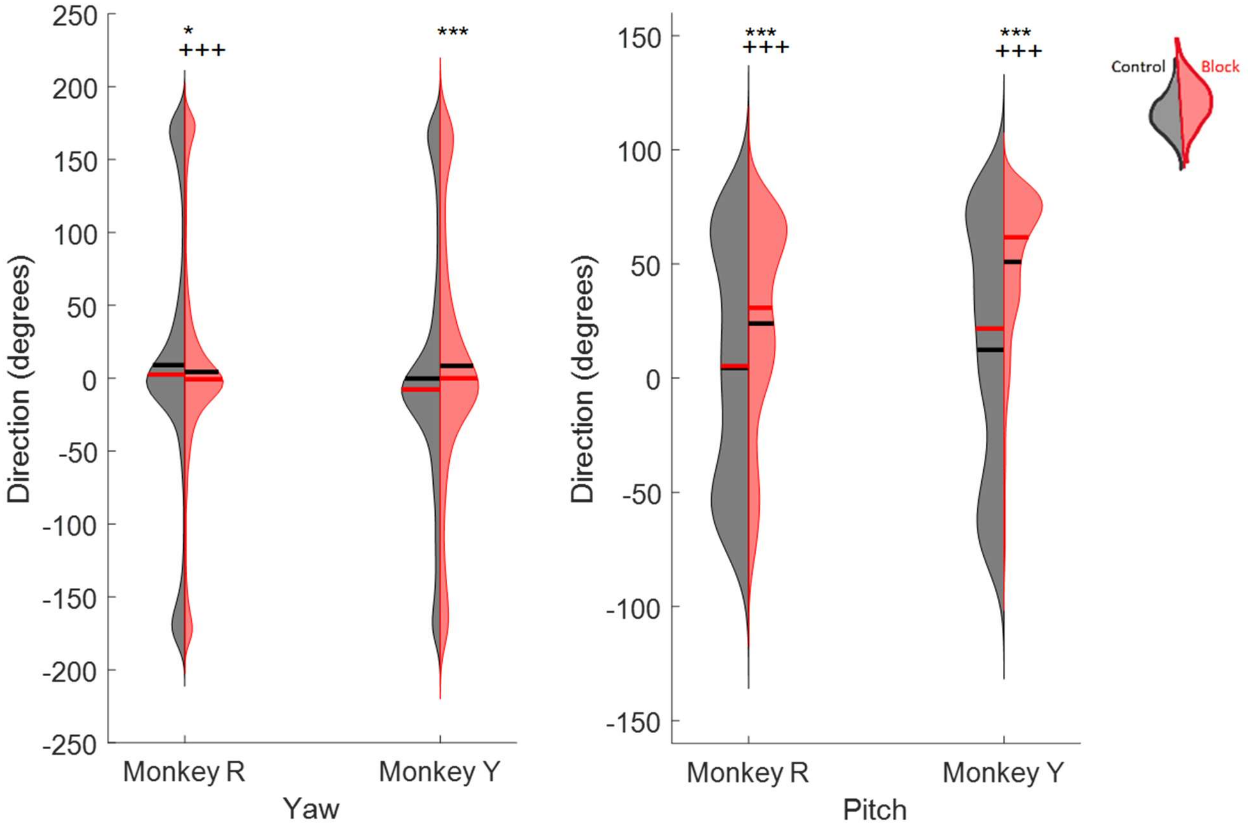


***Supplementary file 2 - Figure 2.*** *Distribution of tongue direction (yaw and pitch) during feeding. Left halves of hemi-violins (black) are control and right halves (red) are nerve block for an individual. Horizontal black lines represent the mean and horizontal red lines the median. Results of two-tailed t-test and f-test are indicated by asterisks and crosses, respectively: *,† $p < 0.05$; **,†† $p < 0.01$; ***,††† $p < 0.001$.*

**Proportion of tuning to yaw and pitch**

Many neurons exhibited modulation of spiking activity to the yaw and pitch angles of tongue movements. Consistent with our 3D angle results, fewer SIo neurons were directionally tuned to yaw and pitch compared to MIo. More neurons were tuned to the pitch direction than the yaw (Chi-square, $p < 0.01$), except in the SIo of Monkey R where the difference was not significant. The changes in the proportion of directionally tuned neurons with the addition of nerve block were also comparable with the changes to 3D directionality. In most cases, there was a decrease with nerve block, except for in the percentage of neurons tuned to yaw in Monkey Y where there was an increase.

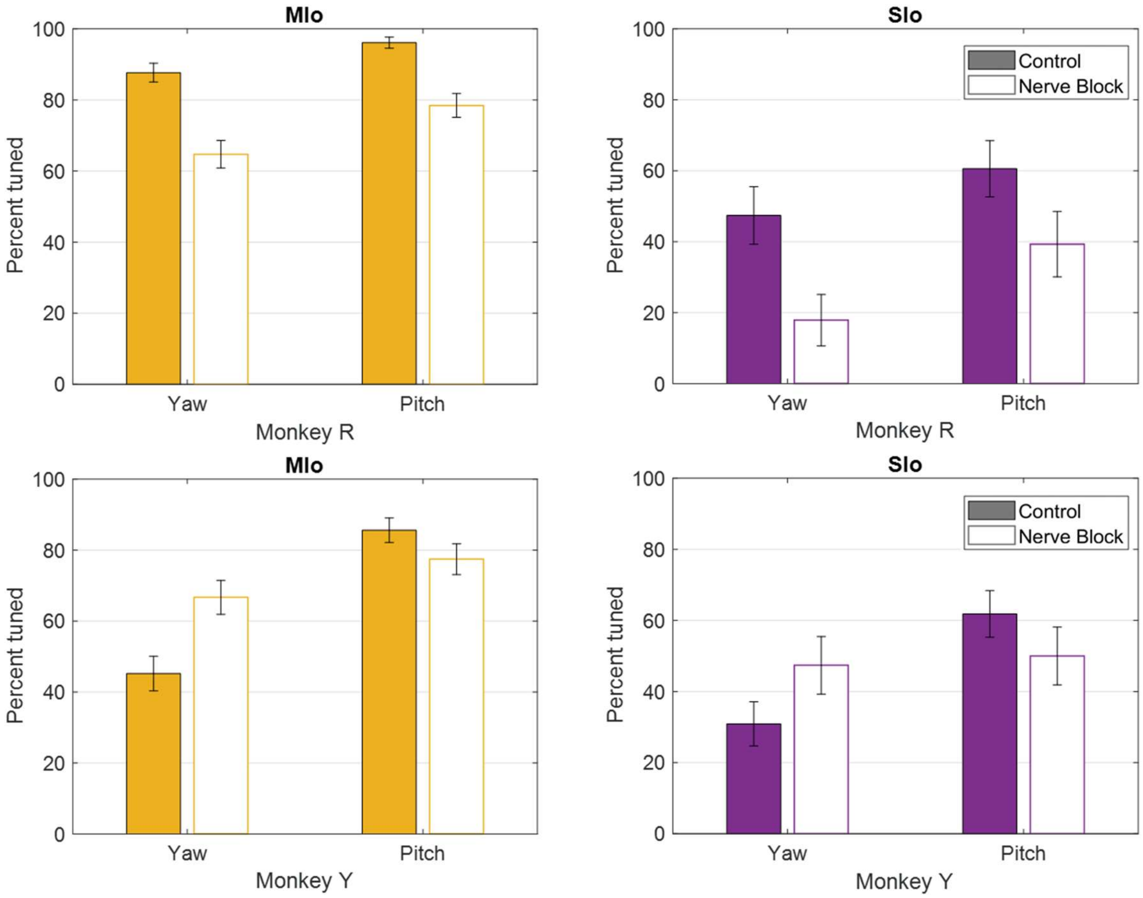


***Supplementary file 2 - Figure 3.*** *Percentage of neurons in MIo (yellow) and SIo (purple) tuned to yaw and pitch under control and nerve block conditions for both subjects. Error bars represent ±1 SE.*

**Distribution of preferred directions**

The distributions of preferred yaw and pitch were comparable between MIo and SIo in both subjects (circular k-test, $p > 0.3$), similar to the 3D angle. Overall, more neurons had an upward preferred pitch than a downward. Whereas in Monkey Y there was a preference across areas for leftward yaw, in Monkey R the PDs were more evenly distributed. Following sensory loss, shifts in mean PD were observed in MIo and SIo of both animals for pitch and/or yaw (circular k-test, $p <$ 0.05).

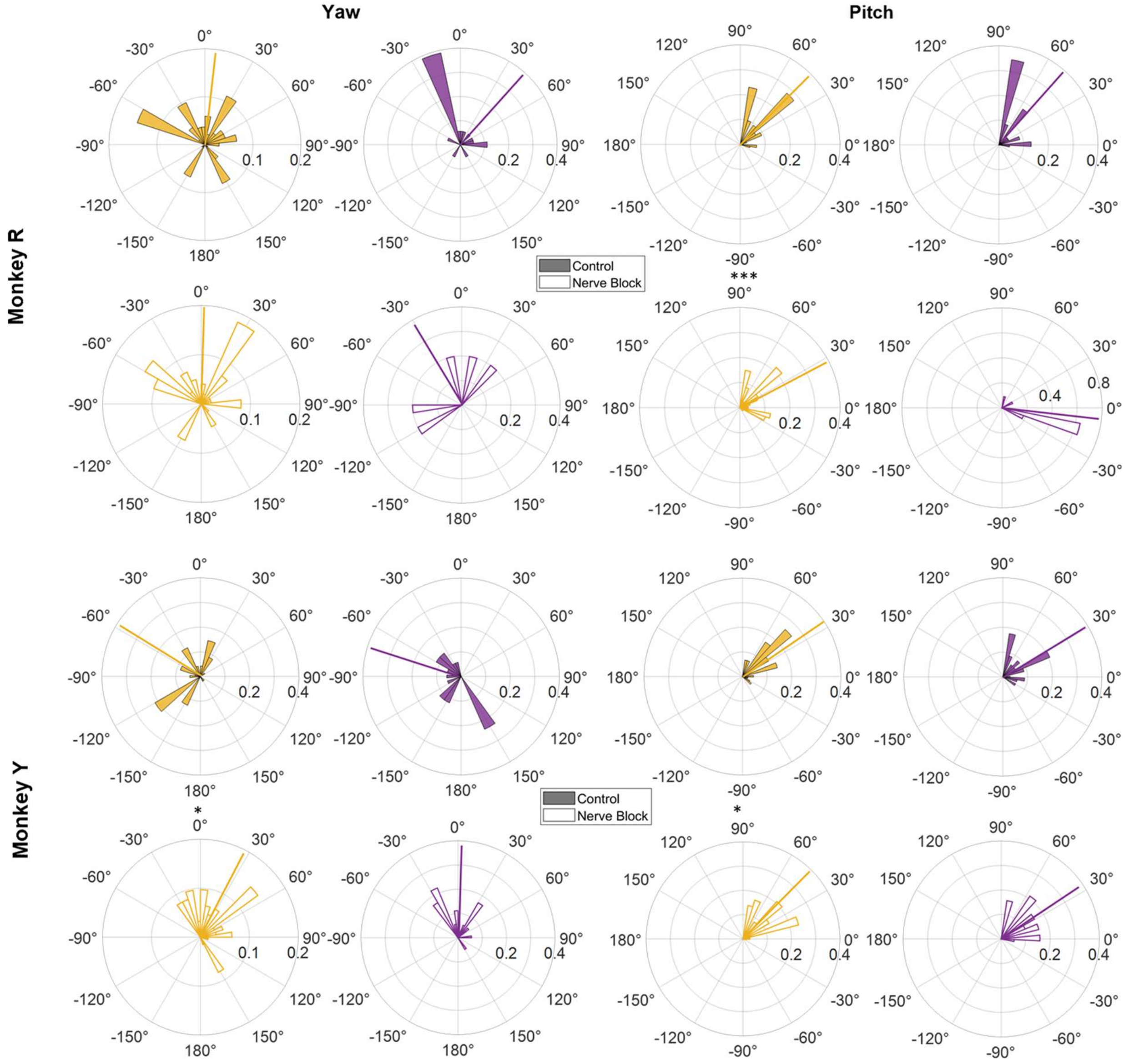


***Supplementary file 2 - Figure 4.*** *Distributions of preferred yaw and pitch in MIo (yellow) and SIo (purple) under control and nerve block conditions. Positive yaw angles represent rightward movement, and positive pitch angles represent upward movement. Polar plots are split into 10° bins with thick colored lines representing the mean PD. Significant circular concentration test (k-test) comparing control and nerve block are indicated by asterisks:* * p < 0.05; ** p < 0.01; *** p < 0.001.
